# Structure-based functional study of a peptide of an ecdysozoan superfamily: reveling a common molecular architecture and receptor-interacting residues

**DOI:** 10.1101/2020.10.29.360867

**Authors:** Yun-Ru Chen, Nai-Wan Hsiao, Shiau-Shan Huang, Chih-Chun Chang, Yi-Zong Lee, Jyuan-Ru Tsai, Hui-Chen Lin, Jean-Yves Toullec, Chi-Ying Lee, Ping-Chiang Lyu

## Abstract

A neuropeptide (Sco-CHH-L), belonging to the crustacean hyperglycemic hormone (CHH) superfamily and preferentially expressed in the pericardial organs (POs) of the mud crab *Scylla olivacea*, was functionally and structurally studied. Its expression levels were significantly higher than the alternative splice form (Sco-CHH) in the POs and increased significantly after animals were subjected to a hypo-osmotic stress. Sco-CHH-L, but not Sco-CHH, significantly stimulated *in vitro* the Na^+^, K^+^-ATPase activity in the posterior (6^th^) gills. Furthermore, solution structure of Sco-CHH-L was resolved using nuclear magnetic resonance spectroscopy revealing that it has an N-terminal tail, three α-helices (α2, Gly^9^−Asn^28^; α3, His^34^−Gly^38^; α5, Glu^62^−Arg^72^), and a π-helix (π4, Cys^43^−Tyr^53^) and is structurally constrained by a pattern of disulfide bonds (Cys^7^-Cys^43^, Cys^23^-Cys^39^, Cys^26^-Cys^52^), which is characteristic of the CHH superfamily-peptides. Sco-CHH-L is topologically most similar to the molt-inhibiting hormone from the Kuruma prawn *Marsupenaeus japonicus* with a backbone root-mean-square-deviation of 3.12 Å. Ten residues of Sco-CHH-L were chosen for alanine-substituted and the resulting mutants were functionally tested using the gill Na^+^, K^+^-ATPase activity assay, showing that the functionally important residues (I2, F3, E45, D69, I71, G73) are located at either end of the sequence, which are sterically close to each other and presumably constitutes the receptor binding sites. Sco-CHH-L was compared with other members of the superfamily revealing a molecular architecture, which is suggested to be common for the crustacean members of the superfamily, with the properties of the residues constituting the presumed receptor binding sites being the major factors dictating the ligand-receptor binding specificity.

## INTRODUCTION

The crustacean hyperglycemic hormone (CHH) peptide superfamily, consisting of a group of structurally and functionally diverse peptides, has long attracted research interest for the study of functional diversification through structural changes (1–10). Peptides constituting the superfamily, once thought to be restricted to crustaceans, have now been clearly shown to be present across clades of Ecdysozoa (11).

Inferences through classical ablation and replacement experiments suggested the presence in the eyestalk of crustaceans of an array of humoral factors – presumptive hormones regulating metabolism, reproduction, molting and growth, cardiac activity, color changes etc. (12) – attesting to the importance of the X-organ/sinus gland (XO/SG) complex, a neuroendocrine tissue in the eyestalk, in the regulation of crustacean physiology. Most of the presumptive factors have since been biochemically purified and characterized, and their biological activity confirmed; these include a group of sequence-related peptides – CHH (the prototypic member of the superfamily), molt-inhibiting hormone (MIH), gonad-inhibiting hormone (GIH) or vitellogenesis-inhibiting hormone (VIH), and mandibular organ-inhibiting hormone (MOIH) (11), that once constituted the CHH family (13–15). Identification in the locust *Schistocerca gregaria* of ion transport peptide (ITP), which stimulates ileal salt and water reabsorption, as an insect member (16) expanded for the first time the existence of the superfamily peptides to other arthropod groups. Studies utilizing approaches of *in silico* data mining further extended its presence to several representative ecdysozoan clades (7,17,18). Recent additions to the superfamily include latrodectin peptides and HAND (helical arthropod-neuropeptide-derived) toxins in the venoms of spiders and centipedes (8,10).

A structural signature of the CHH superfamily is the connectivity of 6 invariant cysteine residues forming a unique pattern of disulfide bonds (C1-C5, C2-C4, and C3-C6). Further, members of the CHH superfamily are placed into one of 2 groups, Type I or Type II, according mainly to the characteristics of the peptide precursors and its mature peptides. A Type-I peptide − CHH, ITP, or their long alternative splice form (CHH-L or ITP-L, see below) – is characterized by having a precursor consisting of a signal peptide, a precursor-related peptide, and a mature peptide, while a Type-II peptide (MIH, VIH, or MOIH) by having a precursor consisting of a signal peptide and a mature peptide, without an intervening precursor-related peptide (11) With regard to the mature peptide, the short-splice form of Type-I peptides (CHH and ITP), but not CHH-L or ITP-L, are C-terminally amidated, which is critically important for the biological activity, while both short- and long-splice forms are N-terminally pyroglutaminated. The majority of Type-II peptides are free at both ends, with some exceptional VIHs and MIHs being C-terminally amidated (11) Analysis of sequence motifs revealed that 2 groups of the peptides (corresponding to the Type-I and Type-II peptides) can be differentiated by group-specific motifs found at the N- and C-termini of the peptides, while the motifs in the center part of the sequence are common to all peptides (1).

CHH-superfamily peptides expressed in crustacean tissues had initially been identified and characterized using respective bioassay and thus named following the biological function the assay is based on. However, it is now widely recognized that these peptides are usually pleotropic in biological activity and in some instances are active in the same assay, *i.e*., functional overlap. For example, while CHH is well established for being involved in metabolism regulation (19–21) and in particular in stress-induced hyperglycemia, it has also been shown to be active in repressing ecdysteroidogenesis in the Y-organ (*i.e*., MIH activity) and methyl farnesoate synthesis in the mandibular organ (*i.e*., MOIH activity) (11). It has been proposed that evolution of the CHH-superfamily peptides involved events of duplication of an ancestral gene, which may have encoded a pleotropic peptide like current CHHs, leading to two main paralogous lineages (Type-I and Type-II peptides). The 2 lineages during the course of evolution may have partitioned the ancestral functions via a process known as subfunctionalization, with the Type-II paralogous lineages evolved to peptides with more specialized functions such as MIHs or VIHs, while Type-I lineages to peptides retaining to a certain degree the functional pleiotropy of the ancestral peptide (7).

Functional and structural polymorphism of the CHH-superfamily is further diversified by a post-transcriptional process – RNA alternative splicing. Thus, for crustacean *chh* gene, there are long- and short-splice forms, which share the same amino acid sequence for the first 40 residues but differ significantly in the rest of the sequences that are encoded by different exons (11). The peptide of long-splice form derived from *chh* gene, the CHH-L peptide, is functionally less characterized than its short form counterpart, CHH. In several species including the mud crab *Scylla olivacea*, where CHH and CHH-L have been functionally tested and compared, CHH-L neither elicited *in vivo* hyperglycemic response nor exerted *in vitro* inhibitory effect on the Y-organ (22–24). The specific functions of CHH-L, which is usually preferentially expressed in non-eyestalk tissues, remain to be characterized.

Studies using recombinant mutants of the CHH-superfamily peptides to pinpoint functionally critical residues have been carried out for MIH of *Marsupenaeus japonicus* (4), CHHs of *S. olivacea* (9), and ITP of *S. gregaria* (2,5), with a general consensus that functionally important residues are located at the N-terminal and C-terminal ends of the peptides. Tertiary structure of 2 crustacean CHH-superfamily peptides, Pej-MIH and Pej-SGP-I-Gly (a glycine-extended precursor of CHH from *M. japonicus*), has been resolved (25,26), as well as that of HAND toxins in the venom of spiders and centipedes (10). Interestingly, despite Pej-MIH and HAND toxins are otherwise structurally similar, structure of the latter peptides lack a C-terminal helix (α5), which is present in the Pej-MIH, a structural difference thought to be related to the functional differences between the 2 types of peptides (10). Moreover, unexpected dissimilarity between the structure of Pej-MIH and Pej-SGP-I-Gly, particularly at the backbone fold at the C-terminus with Pej-SGP-I-Gly lacking an α5, was conspicuously noted (26).

Pericardial organs (POs) are a pair of neurohemal organs, located in the venous cavity surrounding the heart (27), synthesizing and secreting into hemolymph amine and peptide hormones (28,29). One of the physiological processes that are regulated by POs is osmoregulation (28,30). In the present study, Sco-CHH-L, a CHH-L peptide preferentially expressed in the POs of the mud crab *Scylla olivacea* (31) was functionally and structurally characterized. Sco-CHH-L transcript levels were significantly elevated when animals were facing a hypo-osmotic stress. We further report that Sco-CHH-L stimulated *in vitro* the Na^+^/K^+^-ATPase activity in the posterior gills, establishing a specific role for the peptide in the osmoregulation of marine brachyurans. The tertiary structure of Sco-CHH-L was resolved using nuclear magnetic resonance spectroscopy. Functional test of alanine-substituted Sco-CHH-L mutants was performed using the Na^+^, K^+^-ATPase activity assay showing that several functionally critical residues at both ends of the sequence presumably constituting the receptor binding/activation sites. Comparison of Sco-CHH-L with other crustacean peptides, especially the Pej-MIH, suggests a structural architecture likely common to crustacean members of the superfamily.

## RESULTS

### Sco-CHH-L and Sco-CHH transcript and protein expression levels in the pericardial organs

We performed semi-quantitative real-time PCR on the cDNA samples derived from the pericardial organs (POs) with transcript-specific primers to determine the levels of Sco-CHH-L and Sco-CHH transcript; data showed that Sco-CHH-L expressed at levels significantly higher (2.5 folds) than Sco-CHH (Fig. 1A). At the protein levels, an enzyme-linked immunosorbent assay showed that levels of Sco-CHH-L were significantly higher (25.6 folds) than those of Sco-CHH (3.33 vs. 0.13 pmol) (Fig. 1B).

**Figure 1.**
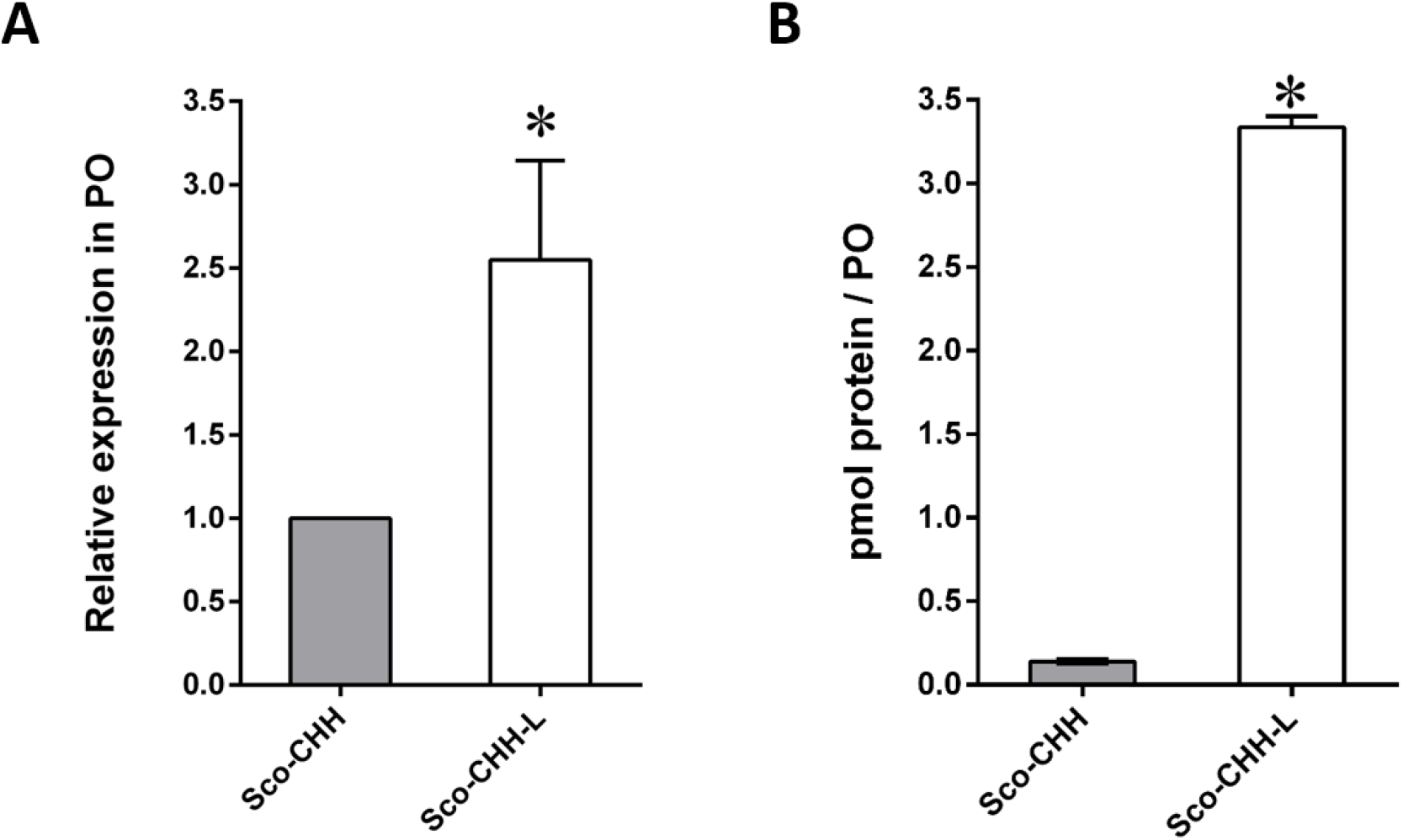
Preferential expression of Sco-CHH-L in the pericardial organs of the mud crab *Scylla olivacea*. (A) Levels of Sco-CHH and Sco-CHH-L transcript determined by a semi-quantitative real-time PCR. Data are means ± SEM. n = 4 for each group. Transcript levels are normalized to a reference gene (*18s rRNA*) and expressed relative to the Sco-CHH levels. (B) Quantification of Sco-CHH and Sco-CHH-L by an ELISA using purified Sco-CHH-L and Sco-CHH as standards. The asterisk (*) indicates significant difference at 5% level.

### Pattern of changes in Sco-CHH-L and Sco-CHH transcript levels under osmotic stresses

Sco-CHH transcript levels in the PO from the 25 ppt-acclimated crabs did not significantly change from the pre-transfer levels (0 hr) when the animals were subjected to either hypo-osmotic (transferred to 5 ppt) or hyper-osmotic (transferred to 45 ppt) stress at any time point examined (Fig. 2). Sco-CHH-L transcript levels in the 25 ppt-acclimated crabs did not significantly change from the pre-transfer levels when subjected to hyper-osmotic stress at any time point, whereas the levels significantly increased in the animals subjected to hypo-osmotic stress at 12 h post-transfer (Fig. 2).

**Figure 2.**
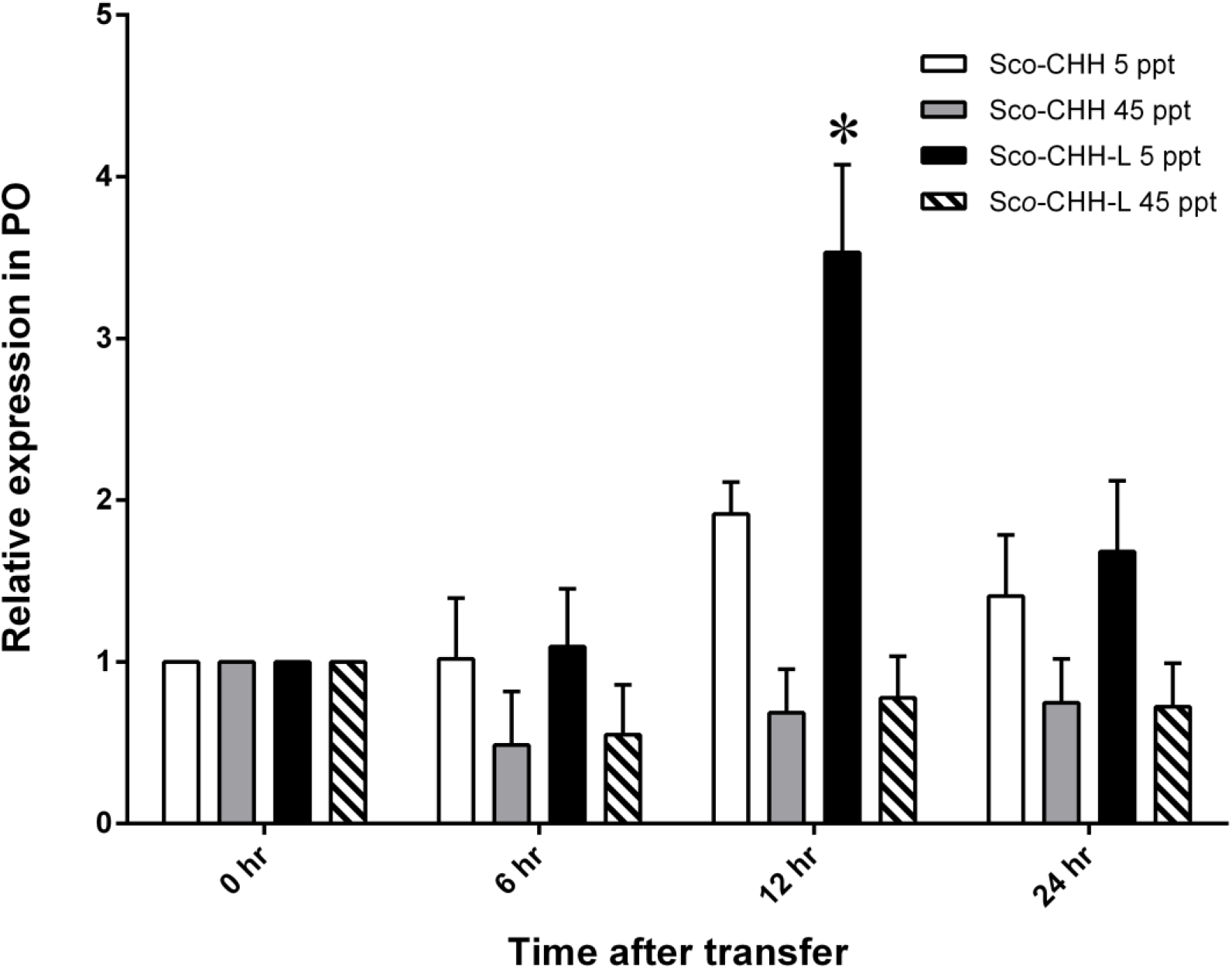
Pattern of changes in Sco-CHH and Sco-CHH-L transcript levels in the pericardial organ of the mud crab *Scylla olivacea* in response to osmotic stresses. Pericardial organs were harvested from 25 ppt-acclimated animals before (0 hr) or at the designated time points (6, 12, 24 hr) after being transfer to a hypo-osmotic (5 ppt) or hyper-osmotic (45 ppt) environment and processed for total RNA extraction and reverse transcription reaction. Transcripts levels were estimated by a semi-quantitative real-time PCR. Data are means ± SEM. n = 4 for each time point. Transcript levels are normalized to a reference gene (*18s rRNA*) and expressed relative to the respective control (0 hr) levels. The asterisk (*) indicates significantly different from at 5% level.

### Characterization of wild-type and mutated Sco-CHH-L peptides

Recombinant proteins were expressed using a bacterial expression system, followed by a refolding reaction, and high performance liquid chromatography (HPLC)-purification. Results of mass spectrometric analysis of the purified recombinant Sco-CHH-L peptides were listed in Table 1, which showed that the observed mass of each peptide is in close agreement with its theoretical value. In addition, circular dichroism (CD) spectrum of each peptide exhibited negative bands at 208 and 222nm (Fig. 3); the calculated α-helical content ranged between 26.2~40.0% (Table 1).

**Table 1.** Identification and characterization of recombinant wild-type and alanine-substituted peptides.

| Peptide | Mass (Da) | | α-helix (%) |
| --- | --- | --- | --- |
|  | Theoretical | Observed |  |
| Sco-CHH | 8527.9 | 8528.0 | 47.0 |
| Sco-CHH-L | 8626.8 | 8626.0 | 40.0 |
| I2A | 8584.7 | 8584.0 | 36.1 |
| F3A | 8550.7 | 8551.0 | 31 |
| V41A | 8598.8 | 8598.3 | 38.5 |
| E45A | 8568.8 | 8568.0 | 38.0 |
| N60A | 8583.8 | 8584.5 | 28.1 |
| E62A | 8568.8 | 8568.5 | 37.5 |
| E63A | 8568.8 | 8568.5 | 37.1 |
| D69A | 8582.8 | 8583.0 | 39.1 |
| I71A | 8584.7 | 8584.0 | 35 |
| G73A | 8640.9 | 8642.0 | 26.2 |

**Figure 3.**
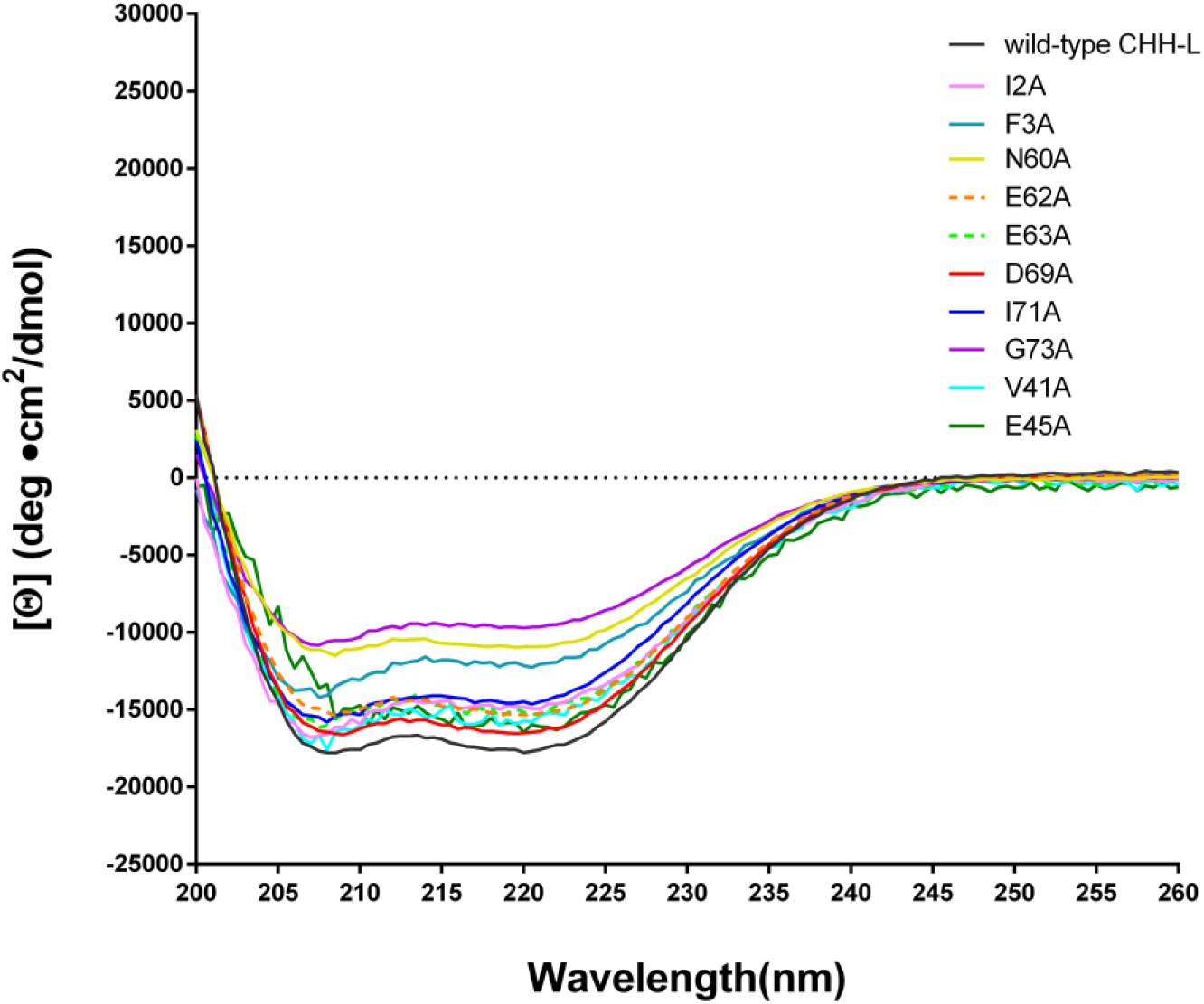
Far-ultra violet circular dichroism spectra of the wild-type rSco-CHH-L and alanine-substituted mutants. CD spectra of the purified peptides (each 15 μM) in phosphate buffered saline (pH 7.4) were recorded at 25°C with a wavelength range of 260 to 200 nm, using a 1-mm path length quartz cell.

### Effects of Sco-CHH and Sco-CHH-L on Na^+^, K^+^-ATPase activity in posterior gills

The wild-type Sco-CHH-L peptide stimulated Na^+^, K^+^-ATPase activity in the posterior gills (gill 6), with the maximal effect being attained at 125 pM (Fig. 4A). On the other hand, Sco-CHH with an amidated C-terminus did not exert any significant effect, even at high doses (500 or 5000 pM) (Fig. 4B).

**Figure 4.**
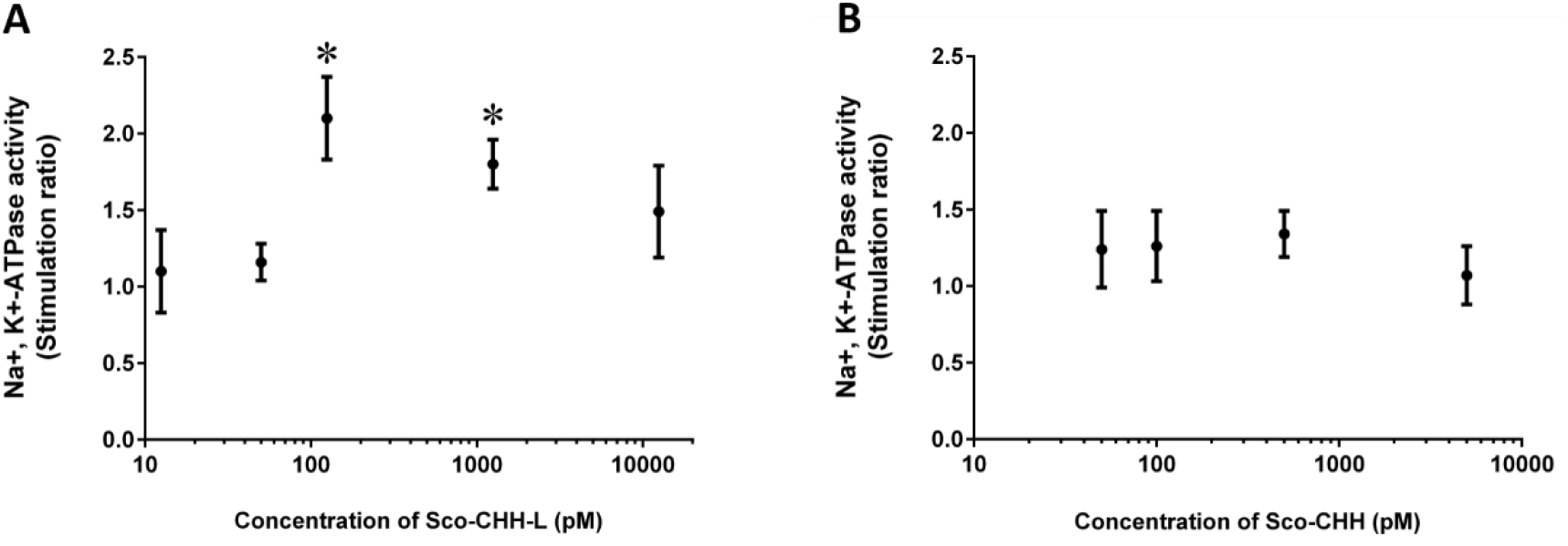
*In vitro* effects of Sco-CHH-L and Sco-CHH on the Na^+^, K^+^-ATPase activity in posterior gills of the mud crab *Scylla olivacea*. The posterior gill (gills 6) were dissected out and received perfusion of (A) Sco-CHH-L (12.5, 50.0, 125.0 and 1250.0 pM) or (B) Sco-CHH (50.0, 100.0, 500.0, and 5000.0 pM). Data are means ± SEM and expressed as the stimulation ratio. n = 4 for each treatment. The asterisk (*) indicates significantly different from vehicle control (saline treatment) at 5% level.

### Solution structure of Sco-CHH-L

We have finished near complete backbone assignments of Sco-CHH-L excepting those for Gln1, Ile2, Phe3, and Asp4 in the N-terminal region. The NMR restraints structural statistics are summarized in Table 2. The initial three dimensional structures of Sco-CHH-L were calculated from a total of 1267 restraints including 1123 NOE-derived distance constraints (338 intra-residues, 337 sequential, 371 medium ranges, and 77 long-range NOEs), 50 hydrogen bond restraints, and 94 dihedral angle restraints (φ: 47, ψ: 47). In the well-defined regions (res. 9-28, 34-38, 43-54, 62-72), the RMSD values of the ensemble conformations (Fig. 5A) were 0.54 ± 0.14 Å for backbone and 1.43 ± 0.27 Å for heavy atoms (Table 2). In the Ramachandran plot for the 10 selected structures, most residues are in the most favored regions (75.4%) or additionally allowed regions (23.2%), with 1.4% of the residues in the generously allowed regions; no residue is in the disallowed regions (Table 2).

**Table 2.** NMR restraints and structural statistics for crustacean hyperglycemic hormone-like peptide (Sco-CHH-L) from *Scylla olivacea* in solution.

|  |  |  |
| --- | --- | --- |
| **Experimental constraints** |  |  |
| NOE |  | 1123 |
| Intra-residue |  | 338 |
| Sequential ( $\vert I-J\vert = 1$ ) | | 337 |
| Medium range ( $\vert I-J\vert \leq 4$ ) | | 371 |
| Long range ( $\vert I-J\vert > 4$ ) | | 77 |
| Hydrogen bond constraints |  | 50 |
| Dihedral angle constraints |  | 94 |
| $\Phi$ | | 47 |
| $\Psi$ | | 47 |
| **RMSD (Å) with respect to the average structure** |  |  |
| Well-defined regions (res. 9-28, 34-38, 43-54, 62-72) |  |  |
| Backbone | | $0.54 \pm 0.14$ |
| Heavy atoms | | $1.43 \pm 0.27$ |
| All residues (res. 1-74) |  |  |
| Backbone | | $1.42 \pm 0.51$ |
| Heavy atoms | | $2.31 \pm 0.51$ |
| **RMSD from idealized covalent geometry** |  |  |
| Bonds (Å) | | $0.0174 \pm 0.0005$ |
| Angles (°) | | $2.1032 \pm 0.0496$ |
| Impropers (°) | | $2.5548 \pm 0.0148$ |
| **Ramachandran statistics** |  |  |
| Most favored regions |  | 75.4% |
| Additionally allowed regions |  | 23.2% |
| Generously allowed regions |  | 1.4% |
| Disallowed regions |  | 0.0% |

**Figure 5.**
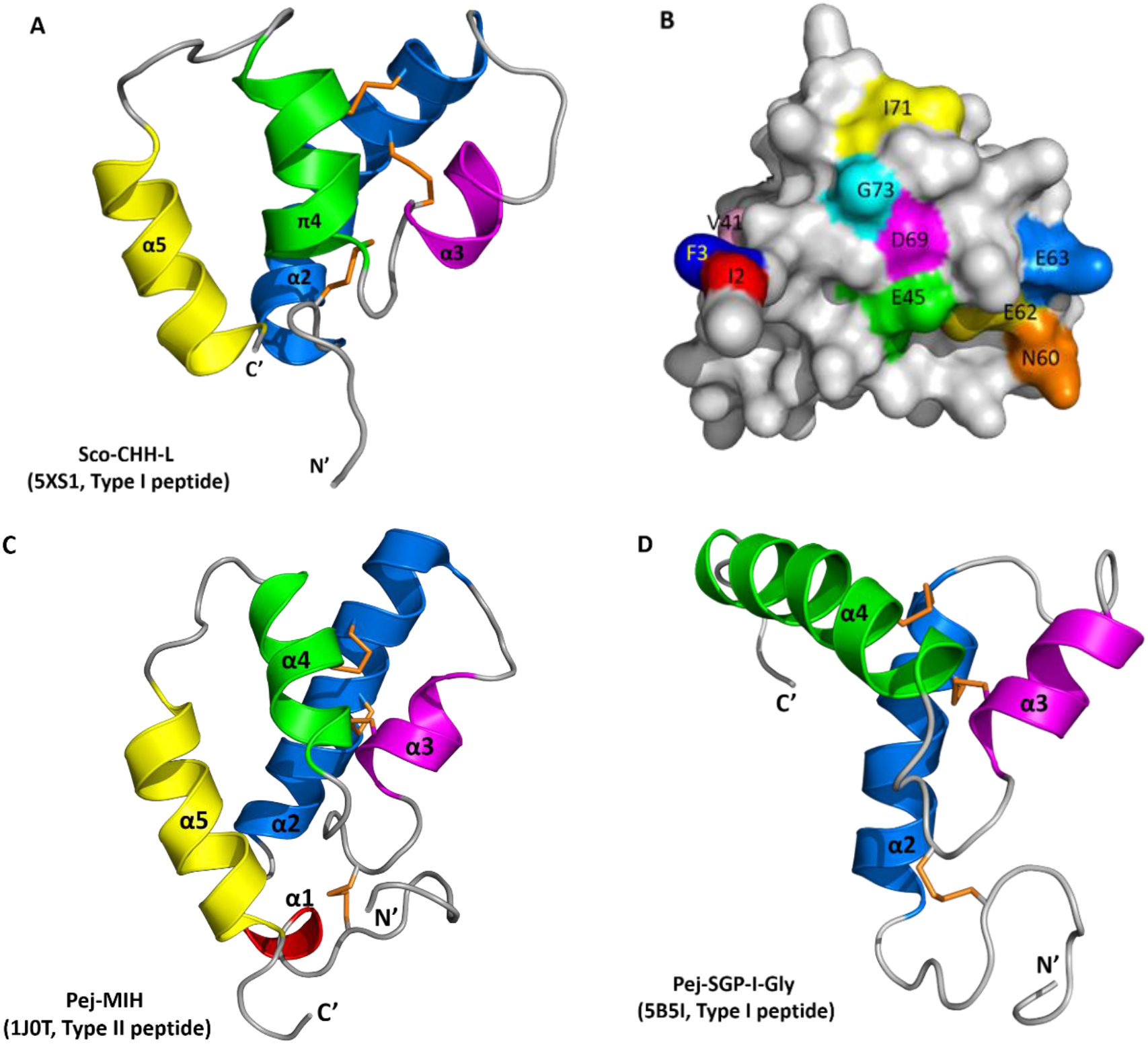
Structure of Sco-CHH-L and its comparison with other CHH/ITP peptides. (A) Ribbon and (B) surface structure of Sco-CHH-L, and ribbon structure of (C) Pej-MIH and (D) Pej-SGP-I-Gly. The helices, N- and C-terminus (N’ and C’) are labeled. (B) Location of the 10 residues that were alanine-substituted in the Sco-CHH-L mutants are labeled on the surface structure. Note that (B) is not viewed from the same angle as (A), but is rotated so that all residues are visible. (A, C, D) Orange sticks indicate disulfide bridging.

The solution structure of Sco-CHH-L contains an N-terminal tail, three α–helices (α2, Gly^9^−Asn^28^; α3, His^34^−Gly^38^; α5, Glu^62^−Arg^72^), a π-helix (π4, Cys^43^−Tyr^53^) between α3 and α5, and three loops between the helices (α2-α3, α3-π4, π4-α5) (Figs. 5 and 6). The first four unassigned residues located in N-terminal loop region are inherently flexible. Sco-CHH-L contains six cysteine residues forming three disulfide bonds (Cys^7^-Cys^43^, Cys^23^-Cys^39^, Cys^26^-Cys^52^), as confirmed by NMR spectrum (Fig. S1), connecting the N-terminal tail and π4, α2 and the loop between α3 and π4, and α2 and π4, respectively. Three aromatic residues in the hydrophobic core of a HAND toxin (Ta1a) important for packing of the structure (Undheim et al., 2015) are conserved in the Sco-CHH-L as Phe^16^, Phe^44^, and Phe^49^ (Fig. 7).

**Figure 6.**
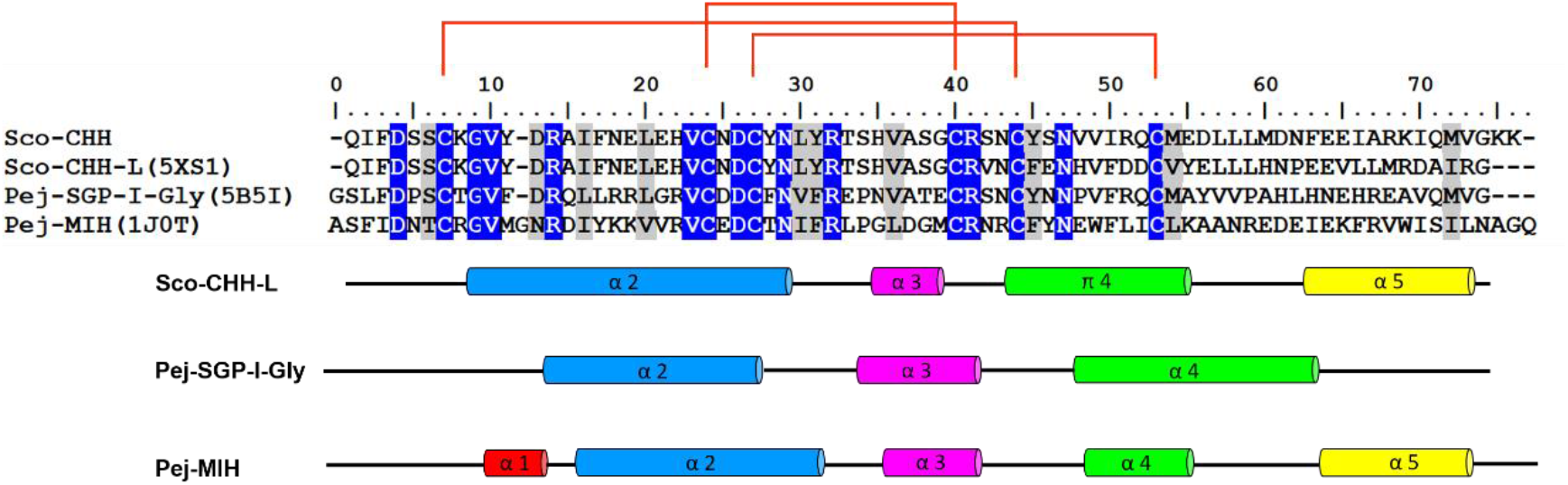
Sequence alignment and secondary structure of 4 CHH/ITP peptides. Amino acid sequences of the mature peptide of 3 Type-I peptides, Sco-CHH, Sco-CHH-L, and Pej-SGP-I-Gly, and that of a Type-II peptide, Pej-MIH, are aligned. The first residue of Sco-CHH sequence is taken as number 1. Accession number for the amino acid sequence is respectively AY372181, EF530127, AB007507, P55847. For Sco-CHH-L, Pej-SGP-I-Gly, and Pej-MIH, location of the secondary structures is given. The conserved pattern of disulfide bridge connectivity (Cys^7^-Cys^43^, Cys^23^-Cys^39^, Cys^26^-Cys^52^) is indicated by orange lines.

**Figure 7.**
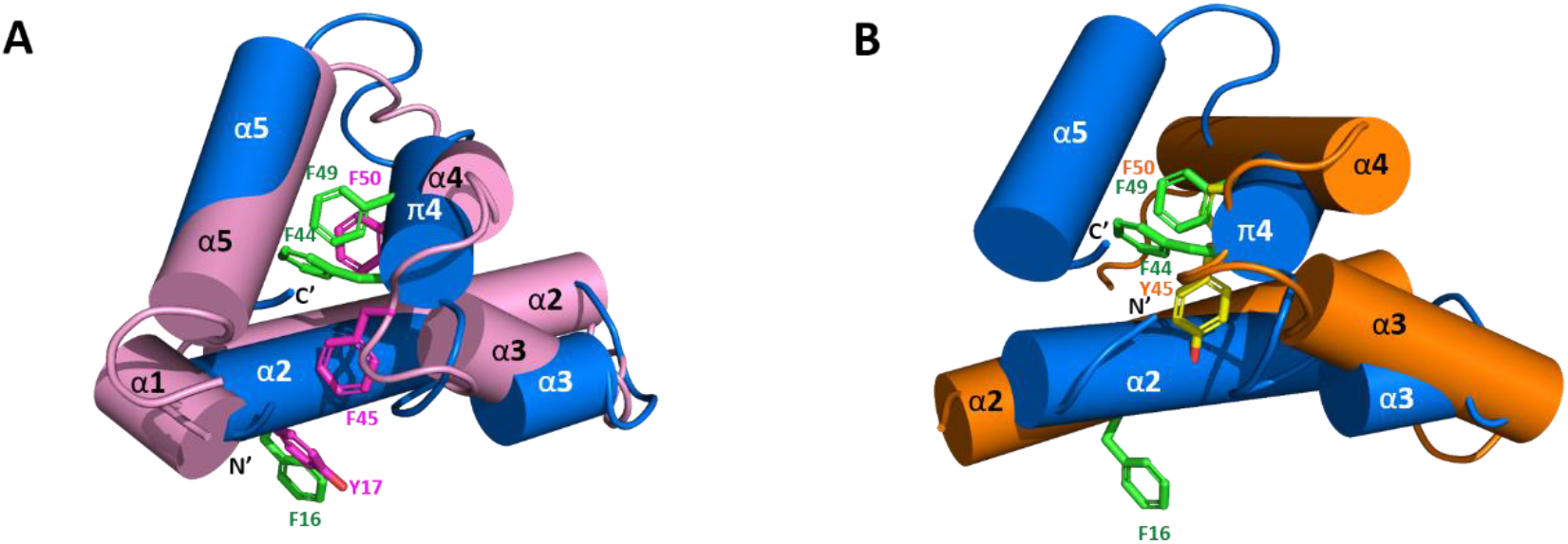
Superposition of Sco-CHH-L with Pej-MIH and Pej-SGP-I-Gly. Structure of Sco-CHH-L (blue) is superposed with (A) Pej-MIH (pink) and with (B) Pej-SGP-I-Gly (orange). Note that Sco-CHH-L is well aligned with Pej-MIH with the N- and C-terminus being brought near together, while the two termini of Pej-MIH are kept away in Pej-SGP-I-Gly. Side chains of the conserved aromatic residues in Sco-CHH-L (Phe^16^, Phe^44^, and Phe^49^ in green), MIH (Tyr^17^, Phe^45^, and Phe^50^ in magenta), and Pej-SGP-I-Gly (Tyr^45^, Phe^50^ in yellow) are shown. Note the aromatic ring of Phe^16^/ Tyr^17^ pointing away from the core structure.

Comparison of the structure of Sco-CHH-L with that of Pej-MIH (Protein Dada Bank: 1J0T), a type II peptide of the CHH superfamily, using Swiss-PdbViewer (Guex and Peitsch, 1997) showed a backbone root mean-square deviation (RMSD) of 3.12 Å, whereas comparison with Pej-SGP-I-Gly (PDB: 5B5I), the precursor of a *Marsupenaeus japonicas* CHH, yielded a RMSD of 4.12 Å.

### Functionally important residues of Sco-CHH-L

To elucidate the residues of Sco-CHH-L important for stimulating the Na^+^, K^+^-ATPase activity in gills 6, ten alanine-substituted Sco-CHH-L peptides (I2A, F3A, V41A, E45A, N60A, E62A, E63A, D69A, I71A, G73A Sco-CHH-L) were produced. The residues chosen for substitution are those with side chains located on the surface of the Sco-CHH-L structure (Fig. 5B). Ile^2^ and Phe^3^ are 2 highly conserved residues in the N-terminus of Type-I peptides and have been shown to be critical for CHH and ITP functions (41–43); Asn^60^ and Asp^69^ were located respectively in loop 3 and α −5 of Sco-CHH-L, positions where the residue property in CHH-L peptides was different to that in CHH peptides (*i.e*., Asn^60^ vs. Asp^60^ and Asp^69^ vs. Ile^69^, Fig. 6; Liu et al., 2015), and alanine substitution of these residues in Sco-CHH significantly decreased its hyperglycemic activity (41); Ile^71^ and Gly^73^, respectively in α-5 and the C-terminal end, are two highly conserved hydrophobic residues among CHH-L peptides (Fig. 6; Liu et al., 2015). In addition, Glu^45^, located in π4, occupies a conserved position where among CHH-L peptides a negatively charged residue (Glu or Asp) is present (Fig. 6; Liu et al., 2015) and 2 negatively residues (Glu^62^ and Glu^63^) located in α5-helix were also chosen for alanine substitution.

When tested in the Na^+^, K^+^-ATPase activity in gills 6, the pump-stimulating activity of 6 mutants (I2A, F3A, E45A, D69A, I71A, G73A Sco-CHH-L) significantly decreased, whereas that of V41A, N60A, E62A, E63A Sco-CHH-L did not, when compared with the wild-type Sco-CHH-L (Fig. 8). The functionally important residues are in steric proximity, despite most are located at either end of the sequence (Fig. 6), which is brought towards each other in the structure of Sco-CHH-L. Thus, Ile^71^, Gly^73^, Glu^45^ and Asp^69^ are on a continuous stretch of the surface, with the 2 N-terminal end residues (Ile^2^ or Phe^3^) being located nearby (Fig. 9).

**Figure 8.**
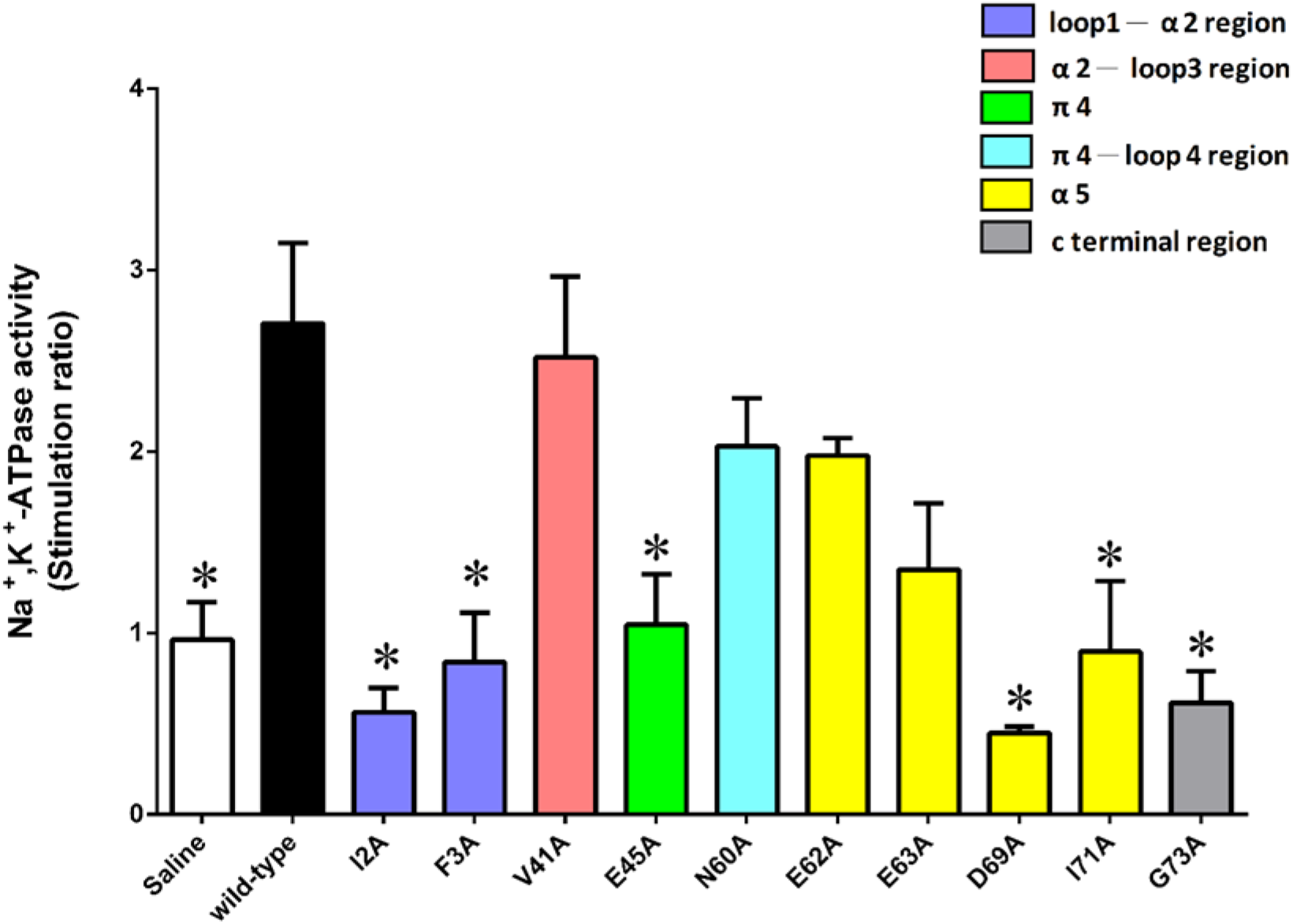
Determination of the functionally important residues of Sco-CHH-L for the Na+, K+-ATPase activity in posterior gills of the mud crab Scylla olivacea. The posterior gill (gills 6) were dissected out and received perfusion of saline, Sco-CHH-L, or an alanine-substituted Sco-CHH-L mutant at 125.0 pM/preparation. Data are means ± SEM and expressed as stimulation ratio. n = 5 for each point. The asterisk (*) indicates significantly different from the Sco-CHH-L group at 5% level.

**Figure 9.**
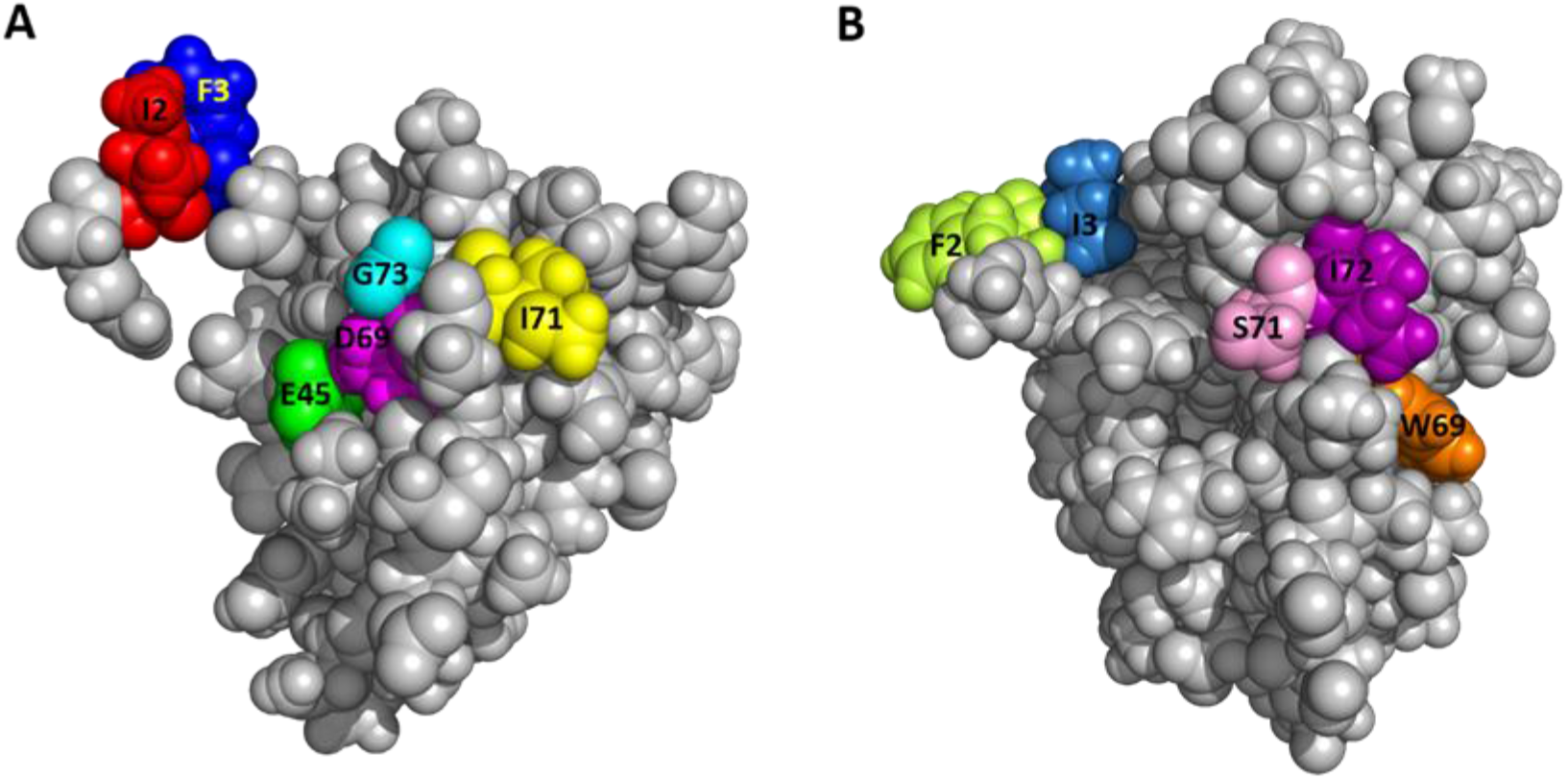
Close proximity of functionally important on the surface of the structure. The space-filling model for (A) Sco-CHH-L and (B) Pej-MIH with the residues possibly constituting receptor binding site are shown in color.

## DISCUSSION

We demonstrated in the present study that a CHH-like peptide (Sco-CHH-L), preferentially expressed in the pericardial organs of the mud crab (*Scylla olivacea*) with its transcript levels being up-regulated by hypo-osmotic stresses, stimulated *in vitro* the Na^+^/K^+^-ATPase activity in the posterior gills, establishing a specific role for the peptide in osmoregulation. Solution structure of Sco-CHH-L was resolved, comparison of which with the structure of other family members defines a molecular topology that is characteristic of the crustacean members of the CHH-superfamily. Pump-activating activity of ten alanine-substituted Sco-CHH-L mutants was tested in the Na^+^, K^+^-ATPase activity assay, showing that the functionally critical residues are present at both ends of the structure, which are sterically close to each other, presumably constituting the receptor binding/activation sites.

One interesting feature of the crustacean peptides belonging to the CHH superfamily is that they are often characterized as being pleotropic in regulatory function, and, in some instances, different peptide members are both active in the same bioassay (although with different degree of potency) (11,32). For example, CHH is best characterized for its role in stress-induced hyperglycemia, but has also been shown to exert inhibitory effect on ecdysteroid synthesis and secretion by the Y-organ, for which MIH is considered the principle regulator (33–36). On the other hand, CHH and CHH-L peptides are alternative splice forms of *chh* gene (11).

CHH-L peptides from several species have been shown possessing neither CHH (hyperglycemic) nor MIH (ecdysteroidogenesis-inhibiting) activity (22–24). In this regard, it is noted that early studies suggested the presence of putative factor(s) in the neuroendocrine tissues, including the eyestalk X-organ/sinus gland complex and pericardial organs, which are involved in osmoregulation (30,37,38). More recent studies have clarified to some extent the identity and molecular target of action of these factors. Thus, CHH, specifically a CHH isoform containing a D-Phe^3^ residue, when *in vivo* injected was able to significantly increase the eyestalk ablation-induced decrease in hemolymph osmolality, Na^+^ concentration, or both, in the freshwater crayfish *Astacus leptodactylus* (39) and the American lobster *Homarus americanus* (40); CHH purified from the crab *Pachygrapsus marmoratus* was able to increase *in vitro* trans-epithelial electrical potential difference and Na^+^ influx in the posterior gills (41); CHH derived from the freshwater crayfish *Cherax quadricarinatus* was effective in restoring stress-induced decrease in hemolymph Na^+^ and K^+^ levels to the pre-stress levels (42). A gut-derived CHH in the crab *Carcinus maenas* was demonstrated to be involved in regulating water and ion uptake during molting, allowing the swelling necessary for successful ecdysis and the subsequent increase in size during postmolt (43). Interestingly enough, apart from the identified CHH, other factors of unknown identity, in the sinus glands were also found to be effective in certain osmoregulatory parameters measured (40,41). In addition, the molecular target(s) on which CHH acts to achieve its osmo-ionic regulation were however not directly addressed by these studies, although it has been suggested that increase in energy availability as a result of CHH-stimulated glycogen mobilization is a probable mode of action (42), a suggestion compatible with the data showing that exposing the shore crabs *C*. *maenas* to dilute seawater increased the levels of glucose in the gills and hemolymph CHH (44). In this regard, it is relevant to note that in the Christmas Island blue crab *Discoplax celeste*, 2 CHHs were found effective in stimulating Na^+^ transport across the gill epithelia in a seasonally dependent manner but have no effect on gill Na^+^/K^+^-ATPase or V-ATPase activity (45). On the other hand, Pt-CHH2, a CHH-L peptide of the crab *Portunus trituberculatus*, was suggested to be a regulator for gill Na^+^/K^+^-ATPase and carbonic anhydrase activity as CHH dsRNA treatment that decreased Pt-CHH2 transcript levels significantly reduced the enzyme activity in the gills (46).

In the present study, we showed that the transcript levels of Sco-CHH-L were increased by exposing animals to a hypo-osmotic stress (25ppt to 5ppt transfer), but not to a hyper-osmotic stress (25ppt to 45ppt transfer). Many marine decapods, including *Scylla* spp., actively regulate hemolymph osmolality in diluted seawater but comply as an osmo-conformer in concentrated seawater (47–49). Thus, the Sco-CHH-L expression pattern in response to osmotic challenges suggested that Sco-CHH-L is involved in osmoregulation when *S. olivace* facing hypo-osmotic stresses. Corroboratively, data showing that Sco-CHH-L peptide, but not Sco-CHH, stimulated *in vitro* the gill Na^+^/K^+^-ATPase indicated that its osmo-regulatory functions are mediated at least in part via stimulating the Na^+^/K^+^-ATPase pump activity. Our data and those mentioned above collectively indicate that CHH and CHH-L peptides are both important factors for regulating water and ionic balance, but these peptides likely do so via acting on distinct molecular targets, perhaps in a concerted manner.

Structure of Sco-CHH-L was compared and contrasted with those of 2 other CHH-superfamily peptides, namely, Pej-MIH and Pej-SGP-I-Gly (25,26). The 6 invariant cysteine residues with a unique disulfide bridge connectivity (C1-C5, C2-C4, C3-C6, Fig. 6) is a signature of the superfamily that shapes the overall fold of the peptides (11). The common core of all the 3 structures is composed of 3 helices, α2, α3, and α4 (π4, in Sco-CHH-L), that have similar topological relationships, with α2 and α3 running in anti-parallel direction and α4 (π4) running orthogonally (Pej-SGP-I-Gly) or obliquely (Pej-MIH and Sco-CHH-L) to α2/α3 (Fig. 5). Three aromatic residues in the hydrophobic core of a venom toxin (Ta1a), a venom peptide of the CHH-superfamily, have been suggested to play a role in the packing of the structure (10). The 3 aromatic residues are conserved in the Pej-MIH and Sco-CHH-L respectively as Tyr^17^, Phe^45^, Phe^50^, and Phe^16^, Phe^44^, and Phe^49^, and only the second (Tyr^44^) and third (Phe^49^) aromatic residues are present in SGP-I-Gly (Fig. 6). It appears that for these peptides only the second and third aromatic residues, being buried in the hydrophobic core, are involved in the packing of the structure, while the side chain of the first conserved residue (in Pej-MIH and Sco-CHH-L) is lying on the surface of the structure (Fig. 7). There are other hydrophobic residues that are conserved across the CHH-superfamily peptides and have been suggested to stabilize the structure (Fig. 6) (11,25); among those residues, only Ile^15^, Val^35^, Val^53^ are located inside the Sco-CHH-L structure and likely contribute to the stabilization of the structure.

Sco-CHH-L shares the most significant topological similarity with Pej-MIH. Besides both having α2, α3, and α4 (or π4), they are the 2 structures that form a topologically equivalent α5 helix, running in an anti-parallel fashion to α4 (or π4), with the C-terminal end being directed towards the N-terminal tail (Figs. 5, 6). Towards the N-terminus, Pej-MIH additionally has a short α1, while the relatively longer α2 in Sco-CHH-L encompasses the region corresponding to that of the α1 in Pej-MIH (Fig. 6). Interestingly, one of the sequence features distinguishing Type I and Type II peptides is that the former group (*e.g*., CHH-L peptide) lacks a Gly^12^ residue, which is present in the latter (*e.g*., MIH) (11). Glycine is known to be a helix breaker, prone to disrupt the regularity of α helical backbone conformation (50). Thus, it is suggested that the presence or absence of the Gly^12^ residue is related to the distinct conformation of the Type I and II peptides in that Sco-CHH-L forms a continuous and long α2, and the corresponding region of Pej-MIH, having a Gly^12^, breaks into 2 helices (α1 and α2), providing a topological signature that differentiates Type I and Type II peptides (Fig. 5), which probably bears functionally important significance as experimental insertion of a Gly^12^ residue into the shrimp CHH (CHH-Gly^12^) significantly decreased its hyperglycemic activity (4).

Superposing Sco-CHH-L on Pej-SGP-I-Gly, an *M. japonicas* CHH precursor, revealed a lower level of resemblance with a higher backbone RMSD. It is somewhat surprising as both structures, as mentioned above, are constrained and shaped by the same disulfide bridge connectivity. The two structure differs significantly in that Pej-SGP-I-Gly, lacking an α5, has a relatively long α4 with the C-terminal tail being kept away from the N-terminal end, while Sco-CHH-L forms, as mentioned above, an α5, which runs in an anti-parallel fashion against α4, with the C-terminal end being close to the N-terminus (Fig. 5). It is interesting to note that HAND toxins, Ta1a and Ssm6a, characteristically lack the C-terminal α5, a structural change thought to be leading to “weaponization” of the ancestral hormonal peptides, by losing the C-terminus of the ancestral sequence that contained α5 (10). Pej-SGP-I-Gly, however, has a rather long C-terminal tail following the α4. As CHH and CHH-L peptides are alternatively spliced forms of *chh* gene having the same sequence up to the first 40 residues but differing significantly after the 40^th^ residues (31), it is possible that the topological differences between Pej-SGP-I-Gly and Sco-CHH-L could be attributed to the differences in the post-40^th^-resiude sequence (Fig. 6). It should also be noted however that Pej-SGP-I-Gly is a non-amidated, glycine-extended precursor of the Pej-SGP-I CHH (26). CHH is post-translationally amidated at the C-terminal end, which is required for full hyperglycemic activity and the α-helical content of CHH increased after the peptide was amidated (24,51,52), indicating that C-terminal amidation renders structural changes. Whether the structure of Pej-SGP-I-Gly will change, particularly at the C-terminus and its topological relationship with the other areas of the structure, as a result of amidation of the peptide, is an interesting issue awaits interrogation.

Effects of alanine-substituted mutant on the gill Na^+^/K^+^-ATPase revealed several residues that are functionally important in the Sco-CHH-L (Fig. 8). Side chains of these residues are located on the surface of the structure, and these residues are located at either end of the peptide but in steric proximity (Fig. 9). It is suggested that the parts of the structure, consisting of these residues, play important roles in forming the binding site for receptor interaction, a suggestion that is applicable to other members of the family, as functionally critical residues at either end of the MIH, CHH, and ITP sequences have also been experimentally demonstrated (2,4,5,9). With respect to MIH, the suggestion is additionally supported by the observation that the 2 ends of the Pej-MIH are, similar to those of Sco-CHH-L, sterically close to each other (Fig. 5), and that the functionally critical residues of each peptide occupy topologically similar positions in the respective structure (Fig. 9). It is further noted that, although the presumed binding sites of the 2 peptides are topologically similar, differences in the side-chain property exist. Thus, whereas both peptides have in common 2 hydrophobic residues, Ile^2^/Phe^3^ in Sco-CHH-L and Phe^2^/Ile^3^ in Pej-MIH, a negatively charged residue (Asp^12^) in the former peptide is replaced by an uncharged residue (Asn^13^) in the latter (Fig. 9). Asp^12^ in Sco-CHH and Asn^13^ in Pej-MIH have been shown to be functionally important (4,9). Similarly, there are 2 negatively charged residue (Glu^45^ and Asp^69^) in Sco-CHH-L that are characteristically absent in Pej-MIH (Fig. 9). It is relevant to mention that receptor binding assay using radiolabeled ligands and membrane preparations derived from potential target tissues suggested the existence of separate binding sites (receptors) for distinct members of the superfamily (53,54). Differences in the property of the residues presumably constituting the binding sites could contribute to the observed receptor binding specificity. Nonetheless, cross binding and activation of receptor by a non-cognate ligand is possible to certain degree because of high topological similarity of the peptides. How the functional specificity and oftentimes the functional overlap as observed for the crustacean members of the CHH-superfamily (see (55)) are achieved through interaction of peptide ligands of a common molecular architecture with their specific receptors represents an endeavor awaits to be pursued for better understanding of functional diversification through structural changes, as well as endocrine regulation of crustacean physiology.

## Experimental procedures

### Animal maintenance

Adult mud crabs *Scylla olivacea* (carapace width: ~7−8 cm) were purchased from Wu-chi Fishing Port, Taichung, Taiwan, transported to the National Changhua University of Education, and acclimated in tanks containing 25 ppt artificial seawater (Blue Treasure), which was continuously aerated and circulated through a bio-filter at a 24–26°C under a 14 h/10 h light/dark cycle (56).

### ELISA for the quantitative analysis of Sco-CHH-L and Sco-CHH in pericardial organs

The pericardial organs were dissected from animals and processed for protein extraction (24). Levels of Sco-CHH-L and Sco-CHH in the extracts were quantified using an enzyme-linked immunosorbent assay (24) with antibodies specifically against Sco-CHH-L or Sco-CHH(31) and purified recombinant Sco-CHH-L and Sco-CHH (24) as standards.

### Gene expression under osmotic stresses

For the experiments of osmotic stress, animals, acclimated to 25 ppt artificial seawater for at least 7 days, were transferred to either 5 ppt or 45 ppt salinity. Crabs were sacrificed at before (0 h), 6 h, 12 h, or 24 h after transfer for harvesting of the pericardial organs, which were frozen directly in liquid nitrogen before use.

Total RNA were isolated from the pericardial organs and reversed transcribed according to Chen et al. (57). A protocol for a semi-quantitative real-time PCR (57) was adopted to quantify Sco-CHH and Sco-CHH-L transcript levels. Primer sequences for amplification of 18s rRNA (reference), Sco-CHH and Sco-CHH-L (targets) genes were listed in Table S1. The real-time PCR amplifications were performed as follows: 95 °C for 10 min followed by 40 cycles of 95 °C for 10 s, 60 °C for 7 s and 72 °C for 7 s. Amplification specificity and efficiency of each pair of primers were experimentally validated according to Chen et al. (57). The comparative threshold cycle method was used to determine transcript levels of the target genes. Thus, levels of Sco-CHH and Sco-CHH-L gene expressions were normalized to those of 18s rRNA gene concurrently amplified. Normalized levels of the target genes at each time point were expressed relative to those at zero hour (arbitrarily designated as the calibrator).

### Construction of Expression Plasmids and Production of Recombinant Peptides

Production, amidation, and characterization of the wild-type recombinant Sco-CHH and Sco-CHH-L have been previously described (24). In addition, construction of the recombinant plasmids each expressing an alanine-substituted mutants of Sco-CHH-L (I2A, F3A, V41A, E45A, N60A, E62A, E63A, D69A, I71A, and G73A Sco-CHH-L) was performed using the QuikChange PCR-mediated Site-Directed Mutagenesis Kit (Agilent) according to the manufacturer's instruction with the plasmid pET-22b(+)-Sco-CHH-L encoding the wild-type Sco-CHH-L (24) as polymerase chain reaction (PCR) template. The synthetic oligonucleotides used in the PCR reactions were listed in Table S1. Each constructed plasmid was sequenced (Mission Biotechnology Inc., Taipei, Taiwan) to ensure the intended sequence was constructed.

Production of recombinant peptides was performed essentially as described in Chang et al. (24) with modifications. Briefly, *Escherichia coli* BL21(DE3) competent cells (Novagen^®^) transformed with each of the plasmids (I2A, F3A, V41A, E45A, N60A, E62A, E63A, D69A, I71A, and G73A pET-22b(+)-Sco-CHH-L) encoding an alanine-substituted Sco-CHH-L. For production of peptides to be used for testing their effects on Na^+^, K^+^-ATPase activity, transformed cells were grown in Luria–Bertani (LB) medium containing ampicillin (100 μg/ml) at 37°C, and the expression was induced with isopropyl β-D-1-thiogalactopyranoside (IPTG) at 1 mM at 37°C for 4 h. For isotope labeling, cells transformed with pET-22b(+)-Sco-CHH-L were grown in M9 minimal medium with ^15^NH_4_Cl (1 g/L) as the nitrogen source and ^13^C-glucose (2 g/L) as the carbon source. Expression of labeled peptides was induced with 0.5 mM IPTG at 25°C for 12 h.

After IPTG induction, cells were harvested by centrifugation (6,000 g, 20 minutes, 4°C), suspended in 10 mM phosphate buffered saline (PBS, pH 7.4), and lysed using a high-pressure homogenizer (GW technologies, Taiwan). After centrifugation (30,000 g, 30 minutes, 4°C), both supernatants and pellets were analyzed using 16.5% Tricine–Sodium Dodecyl Sulphate–Polyacrylamide Gel Electrophoresis (Tricine– SDS–PAGE) (58), which showed most expressed proteins were in the pellets.

A refolding protocol (24) with modification was used to refold the recombinant peptides in the pellets. Briefly, the pellets were first denatured in 100 mM Tris-HCl, at pH 8.0, containing 6M guanidine-HCl and the reactions incubated at 4°C overnight (12 h ~16 h) with stirring. After centrifugation (30,000 g, 30 minutes, 4°C), the supernatant was applied to a C_18_ Sep-Pak^®^ cartridge (WAT043345, Waters) and eluted with 60% acetonitrile. The collected fractions were lyophilized and the lyophilized samples dissolved in 250 mM Tris-base buffer with 8 M Urea at pH 8.5 (1:100 w/v). Subsequently, the dissolved samples were slowly diluted with 19 volumes of a refolding buffer containing 250 mM Tris-base, 10% glycerol, 5 mM cysteine, 0.5 mM cystine, pH 8.3, after which urea was added to the refolding reaction to a final concentration of 1.5 M. The refolding reaction was incubated 4°C for 20 h with stirring (300 rpm). Subsequently, the refolding reaction was filtered through a PES membrane (0.22 μm) and the filtrate concentrated using a ultrafiltration apparatus with a 5 kDa molecular mass cut-off (Vivaflow^®^ 200 cassette, Sartorius Stedim Biotech). The concentrate was separated by reversed phase (RP-HPLC) system with water/acetonitrile gradient to obtain purified recombinant peptides (24).

Mass and purity of the purified recombinant peptides were determined using Micro Mass Quattro Ultima mass spectrometer with the electron spray ionization (ESI-MASS). Concentration of the recombinant peptides was determined using Pierce BCA Protein Assay Kit (Thermo Fisher Scientific, USA) with bovine serum albumin as the standard.

### Circular dichroism spectral analysis of recombinant wild-type and alanine-substituted Sco-CHH-L

Far-ultraviolet circular dichroism (CD) spectra of the recombinant peptides, that were used in the gill Na^+^, K^+^-ATPase activity assay, were measured by an Aviv CD spectrometer (Model 202, Aviv Biomedical Inc., Lakewood, NJ) in phosphate buffered saline (150 mM NaCl, 8mM Na_2_HPO_3_, 2mM KH_2_PO_4_, pH 7.4) with 15 μM of peptides. An aliquot (200 μl) of the samples was placed in a 1-mm cell path length quartz cell and each spectrum, an average of three scans, was collected at 25°C with a wavelength range of 260 to 200 nm in 0.5 nm increments. Raw data were baseline corrected, smoothed, and transformed to obtain spectra in units of mean residue ellipticity according to Zhou et al. (59). The ellipticity at 222 nm was used to estimate the α-helical content of the peptides using DichroWeb sever (60).

### Effects of wild-type peptides and mutated Sco-CHH-L peptides on Na^+^, K^+^-ATPase activity

To establish a dose-dependent curve for the effect of Sco-CHH-L and Sco-CHH on the Na^+^, K^+^-ATPase activity in the 6^th^ pair of gills (gills 6), the enzyme-specific activity was determined using an assay previously developed for *S*. *paramamosain* (56) with some modifications. After being acclimated for 7 days in 25 ppt seawater, crabs were sacrificed on ice and gills 6 were removed from the branchial chambers. Hemolymph of the tissues was flushed out using excessive volume of Pantin’s saline injected through the cut end of the efferent and afferent vessels using a hypodermic syringe with a 25-G needle. Subsequently, one of the gills 6 is perfused with Pantin’s saline, while the contralateral one with saline containing the desired dose of wild-type Sco-CHH-L or Sco-CHH. Preliminary test indicated that there was no significant difference in basal enzyme activity between the left and right gills (left: 19.09 + 3.38 μmole Pi/mg protein/hr; right: 19.59 + 2.34 μmole Pi/mg protein/hr, n = 3 for each group), hence, gills were randomly assigned to the saline-treated or peptide-treated groups. Tissues were then individually incubated at 25°C for 30 min in culture dish with saline or saline containing the recombinant peptides at the same dose that was used to perfuse the tissues. After incubation, each gill was cut into small pieces and homogenized in 500 μl of homogenization buffer (25 mM Tris-HCl, 250 mM sucrose, 20 mM EDTA, 0.4% sodium deoxycholate, pH 7.4) containing protease inhibitors (in final concentration 16.5 μM antipain, 10.8 μM leupeptin, 320 μM benzamidine, and 50 μM aprotinin, Sigma) using an ultrasonic processor. Crude homogenates were centrifuged (6,000 g, 4°C, 15 min) and the resulting supernatants re-centrifuged (30,000 g, 4°C, 20 min). An aliquot of the final supernatants was saved for determination for total protein by Bio-Rad Protein Assay Kit (#5000002, Bio-Rad, USA) with bovine serum albumin as the standard at 595 nm, and the rest of the supernatants immediately used for determination of Na^+^, K^+^-ATPase activity assay.

The protocol for Na^+^, K^+^-ATPase activity has been described by Chung and Lin (56). The Na^+^, K^+^-ATPase activity was assayed by adding 10 μl of the supernatant to 0.4 ml of reaction buffer, which was, for the ouabain-free group, 20 mM imidazole, 100 mM NaCl, 30 mM KCl, and 10 mM MgCl_2_ at pH 7.4, or for the ouabain group, 20 mM imidazole, 130 mM NaCl, 10 mM MgCl_2_, and 1 mM ouabain at pH 7.4. Reaction was initiated by adding 0.1 ml of the ATP stock solution (25mM Na_2_ATP), incubated for 15 min at 30°C, quenched by adding 0.2 ml of an ice-cold trichloroacetic acid solution (30% w/v), and centrifuged (1,640 g, 4°C,10 min). An aliquot (0.5 ml) of the supernatants was taken and subsequently reacted with 1 ml ice-cold Bonting’s colorimetric solution (176 mM FeSO_4_, 560 mM H_2_SO_4_, 8.1 mM ammonium molybdate) at 20°C water bath for 20 min. Concentration of the inorganic phosphate (Pi) was measured spectrophotometrically at 700 nm (Hitachi, Japan) according to Peterson's method (61). The Na^+^, K^+^-ATPase activity was calculated as the difference in the levels of Pi between the ouabain group and the ouabain-free group and normalized by tissue total proteins (56,62,63).

The effect of peptides on Na^+^, K^+^-ATPase activity was calculated and expressed as the stimulation ratio, i.e., the activity of the peptide-treated gill over the enzyme-specific activity of the contralateral saline-treated gill.

To test the effect of rSco-CHH-L mutants on the Na^+^, K^+^-ATPase activity, gills 6 were dissected, treated, and incubated with saline, wild-type Sco-CHH-L, or each of the Sco-CHH-L mutants (peptide concentration at 125 pM), processed for determination of the enzyme-specific activity, and calculated for the stimulation ratio as described above.

### Nuclear Magnetic Resonance spectroscopy and structure determination

It has been reported that Pej-MIH did not completely dissolve at high concentrations (~1 mM) in water or PBS buffer (25). Similarly, pilot tests showed that rSco-CHH-L was quite insoluble at concentration higher than 0.3 mM over a range of pH (pH 4.0 – 7.4). Thus, rSco-CHH-L was dissolved (15 μM) in 10 mM citric acid-sodium phosphate (pH 3.0, 4.0, or 5.0) and measured for CD spectra. The results shown in Figure S2 revealed that Sco-CHH-L at pH 3.0 – 5.0 exhibited negative signal at 208 and 222 nm, with an estimated α-helical content of 42%, which is close to the content measured for Sco-CHH-L in PBS (40%) at pH 7.4. To further improve solubility for NMR experiments, we then added L-Glutamate, which has been shown to facilitate protein solubility and long-term stability (64), to the peptide solution in acidic pH range buffer. The results showed that rSco-CHH-L could be dissolved over 1 mM in 10 mM citric acid-sodium phosphate buffer (100 mM sodium chloride and 10 mM potassium chloride) at pH 3.0 with 50 mM L-Glutamate. Heteronuclear single quantum coherence (HSQC) spectroscopy analysis of the dissolved peptides indicated that HSQC cross peaks were well dispersed (Fig. S3). Sco-CHH-L thus dissolved was used for subsequent NMR experiments.

NMR data were acquired on a Bruker Avance 600 MHz, 850 MHz and Varian 700 MHz spectrometer. For structure determination, rSco-CHH-L (^15^N-labelled or ^13^C/^15^N-double-labelled) was dissolved at 0.9 mM in buffer consisting of 10 mM citrate phosphate, 100 mM sodium chloride, 10 mM potassium chloride, 50 mM L-Glutamate and 0.02% sodium azide and 10 % (v/v) D_2_O at pH 3.0, and subjected to NMR experiments at 25°C. Samples also contained 4,4-dimethyl-4-silapentane-1-sulfonic acid (DSS) as an internal standard (^1^H = 0 ppm). Backbone assignment was accomplished with HNCA, HN(CO)CA, HNCACB, CBCA(CO)NH, HNCO and HN(CA)CO experiments. The chemical-shift resonance assignments of remaining atoms were accomplished using both ^1^H–^15^N HSQC–NOESY with the assistance of through-bond correlation spectra. To obtain side chain resonances, combined information from 3D ^1^H–^15^N HSQC– TOCSY, HBHA(CO)NH, H(CC)(CO)NH, CC(CO)NH, HCCH-COSY and HCCH-TOCSY spectra were analyzed. The NMR spectra were processed with Varian Vnmrj or NMRpipe and the resonance assignments for the set of spectra were analyzed using SPARKY v3.114 (65). The assigned resonances were deposited into the Biological Magnetic Resonance Bank (BMRB) under the entry number BMRB36099.

Structural calculations were carried out with the standard simulated annealing protocol using the CNS program(66). Several rounds of calculations were used to eliminate the ambiguity in the assignment. The 10 lowest energy structures selected out of 200 calculated structures were then refined by the explicit water model using the GROMOS force fields (67). The final structures were evaluated with PROCHECK-nmr software (68). The coordinates of the ensemble of structures have been deposited into the Protein Data Bank (PDB) as entry 5XS1.

### Data and statistical analysis

Data are expressed as mean values ± S.E.M. Student’s *t*-test or one-way ANOVA (with post hoc Tukey’s pairwise comparison) were employed for statistical analysis (SigmaStat v. 3.5).

## Data availability

The atomic coordinates, structure factors, and chemical shift assignments (PDB ID 5XS1) have been deposited in the Protein Data Bank (http://wwpdb.org/). The rest of the data are contained within the manuscript.

## Acknowledgments

The authors thank the NMR Instrumentation Center at National Tsing Hua University, Hsinchu, and Academia Sinica High-Field NMR Center (HFNMRC) for technical support. HFNMRC is funded by Academia Sinica Core Facility and Innovative Instrument Project (AS-CFII-108-112).

## REFERENCE

1. Lacombe, C., Greve, P., and Martin, G. (1999) Overview on the sub-grouping of the crustacean hyperglycemic hormone family. Neuropeptides 33, 71–80

2. Wang, Y. J., Zhao, Y., Meredith, J., Phillips, J. E., Theilmann, D. A., and Brock, H. W. (2000) Mutational analysis of the C-terminus in ion transport peptide (ITP) expressed in Drosophila Kc1 cells. Arch Insect Biochem Physiol 45, 129–138

3. Chan, S. M., Gu, P. L., Chu, K. H., and Tobe, S. S. (2003) Crustacean neuropeptide genes of the CHH/MIH/GIH family: implications from molecular studies. Gen Comp Endocrinol 134, 214–219

4. Katayama, H., Ohira, T., Nagata, S., and Nagasawa, H. (2004) Structure-activity relationship of crustacean molt-inhibiting hormone from the kuruma prawn Marsupenaeus japonicus. Biochemistry 43, 9629–9635

5. Zhao, Y., Meredith, J., Brock, H. W., and Phillips, J. E. (2005) Mutational analysis of the N-terminus in Schistocerca gregaria ion-transport peptide expressed in Drosophila Kc1 cells. Arch Insect Biochem Physiol 58, 27–38

6. Padhi, A., Verghese, B., Otta, S. K., Varghese, B., and Ramu, K. (2007) Positive Darwinian selection on crustacean hyperglycemic hormone (CHH) of the green shore crab, Carcinus maenas. In Silico Biol 7, 355–367

7. Montagné, N., Desdevises, Y., Soyez, D., and Toullec, J.-Y. (2010) Molecular evolution of the crustacean hyperglycemic hormone family in ecdysozoans. BMC Evolutionary Biology 10, 62

8. McCowan, C., and Garb, J. E. (2014) Recruitment and diversification of an ecdysozoan family of neuropeptide hormones for black widow spider venom expression. Gene 536, 366–375

9. Liu, C. J., Huang, S. S., Toullec, J. Y., Chang, C. Y., Chen, Y. R., Huang, W. S., and Lee, C. Y. (2015) Functional Assessment of Residues in the Amino- and Carboxyl-Termini of Crustacean Hyperglycemic Hormone (CHH) in the Mud Crab Scylla olivacea Using Point-Mutated Peptides. PLoS One 10, e0134983

10. Undheim, E. A., Grimm, L. L., Low, C. F., Morgenstern, D., Herzig, V., Zobel-Thropp, P., Pineda, S. S., Habib, R., Dziemborowicz, S., Fry, B. G., Nicholson, G. M., Binford, G. J., Mobli, M., and King, G. F. (2015) Weaponization of a Hormone: Convergent Recruitment of Hyperglycemic Hormone into the Venom of Arthropod Predators. Structure 23, 1283–1292

11. Webster, S. G., Keller, R., and Dircksen, H. (2012) The CHH-superfamily of multifunctional peptide hormones controlling crustacean metabolism, osmoregulation, moulting, and reproduction. Gen Comp Endocrinol 175, 217–233

12. Kleinholz, L. H. (1976) Crustacean Neurosecretory Hormones and Physiological Specificity. American Zoologist 16, 151–166

13. Kegel, G., Reichwein, B., Tensen, C. P., and Keller, R. (1991) Amino acid sequence of crustacean hyperglycemic hormone (CHH) from the crayfish, Orconectes limosus: emergence of a novel neuropeptide family. Peptides 12, 909–913

14. Keller, R. (1992) Crustacean neuropeptides: structures, functions and comparative aspects. Experientia 48, 439–448

15. Soyez, D. (1997) Occurrence and diversity of neuropeptides from the crustacean hyperglycemic hormone family in arthropods. A short review. Ann N Y Acad Sci 814, 319–323

16. Meredith, J., Ring, M., Macins, A., Marschall, J., Cheng, N. N., Theilmann, D., Brock, H. W., and Phillips, J. E. (1996) Locust ion transport peptide (ITP): primary structure, cDNA and expression in a baculovirus system. J Exp Biol 199, 1053–1061

17. Christie, A. E. (2008) Neuropeptide discovery in Ixodoidea: an in silico investigation using publicly accessible expressed sequence tags. Gen Comp Endocrinol 157, 174–185

18. Christie, A. E., Nolan, D. H., Garcia, Z. A., McCoole, M. D., Harmon, S. M., Congdon-Jones, B., Ohno, P., Hartline, N., Congdon, C. B., Baer, K. N., and Lenz, P. H. (2011) Bioinformatic prediction of arthropod/nematode-like peptides in non-arthropod, non-nematode members of the Ecdysozoa. Gen Comp Endocrinol 170, 480–486

19. Santos, E. A., and Keller, R. (1993) Crustacean hyperglycemic hormone (CHH) and the regulation of carbohydrate metabolism: Current perspectives. Comparative Biochemistry and Physiology Part A: Physiology 106, 405–411

20. Li, W., Chiu, K. H., Tien, Y. C., Tsai, S. F., Shih, L. J., Lee, C. H., Toullec, J. Y., and Lee, C. Y. (2017) Differential effects of silencing crustacean hyperglycemic hormone gene expression on the metabolic profiles of the muscle and hepatopancreas in the crayfish Procambarus clarkii. PLoS One 12, e0172557

21. Li, W., Chiu, K. H., and Lee, C. Y. (2019) Regulation of amino acid and nucleotide metabolism by crustacean hyperglycemic hormone in the muscle and hepatopancreas of the crayfish Procambarus clarkia. PLoS One 14, e0221745

22. Dircksen, H., Bocking, D., Heyn, U., Mandel, C., Chung, J. S., Baggerman, G., Verhaert, P., Daufeldt, S., Plosch, T., Jaros, P. P., Waelkens, E., Keller, R., and Webster, S. G. (2001) Crustacean hyperglycaemic hormone (CHH)-like peptides and CHH-precursor-related peptides from pericardial organ neurosecretory cells in the shore crab, Carcinus maenas, are putatively spliced and modified products of multiple genes. Biochem J 356, 159–170

23. Ohira, T., Tsutsui, N., Nagasawa, H., and Wilder, M. N. (2006) Preparation of two recombinant crustacean hyperglycemic hormones from the giant freshwater prawn, Macrobrachium rosenbergii, and their hyperglycemic activities. Zoological science 23, 383–391

24. Chang, C. C., Tsai, K. W., Hsiao, N. W., Chang, C. Y., Lin, C. L., Watson, R. D., and Lee, C. Y. (2010) Structural and functional comparisons and production of recombinant crustacean hyperglycemic hormone (CHH) and CHH-like peptides from the mud crab Scylla olivacea. Gen Comp Endocrinol 167, 68–76

25. Katayama, H., Nagata, K., Ohira, T., Yumoto, F., Tanokura, M., and Nagasawa, H. (2003) The solution structure of molt-inhibiting hormone from the Kuruma prawn Marsupenaeus japonicus. J Biol Chem 278, 9620–9623

26. Tsutsui, N., Sakamoto, T., Arisaka, F., Tanokura, M., Nagasawa, H., and Nagata, K. (2016) Crystal structure of a crustacean hyperglycemic hormone (CHH) precursor suggests structural variety in the C-terminal regions of CHH superfamily members. FEBS J 283, 4325–4339

27. Carlisle, D. B., and Knowles, F. G. W. (1953) Neurohæ mal Organs in Crustaceans. Nature 172, 404–405

28. Cooke, I., and Sullivan, R. (1982) Hormones and neurosecretion. In “The Biology of Crustacea Vol. 3” Ed by Atwood HL, Sandeman DC. Academic Press, New York

29. Christie, A. E., Skiebe, P., and Marder, E. (1995) Matrix of neuromodulators in neurosecretory structures of the crab Cancer borealis. J Exp Biol 198, 2431–2439

30. Kamemoto, F. I. (1976) Neuroendocrinology of Osmoregulation in Decapod Crustacea. American Zoologist 16, 141–150

31. Tsai, K. W., Chang, S. J., Wu, H. J., Shih, H. Y., Chen, C. H., and Lee, C. Y. (2008) Molecular cloning and differential expression pattern of two structural variants of the crustacean hyperglycemic hormone family from the mud crab Scylla olivacea. Gen Comp Endocrinol 159, 16–25

32. Lee, C.-Y., Tsai, K.-W., Tsai, W.-S., Jiang, J.-Y., and Chen, Y.-J. (2014) Crustacean hyperglycemic hormone: structural variants, physiological function, and cellular mechanism of action. Journal of Marine Science and Technology 22, 75–81

33. Yasuda, A., Yasuda, Y., Fujita, T., and Naya, Y. (1994) Characterization of crustacean hyperglycemic hormone from the crayfish (Procambarus clarkii): multiplicity of molecular forms by stereoinversion and diverse functions. Gen Comp Endocrinol 95, 387–398

34. Chung, J. S., and Webster, S. G. (2003) Moult cycle-related changes in biological activity of moult-inhibiting hormone (MIH) and crustacean hyperglycaemic hormone (CHH) in the crab, Carcinus maenas. From target to transcript. Eur J Biochem 270, 3280–3288

35. Chang, E. S., Prestwich, G. D., and Bruce, M. J. (1990) Amino acid sequence of a peptide with both molt-inhibiting and hyperglycemic activities in the lobster, Homarus americanus. Biochem Biophys Res Commun 171, 818–826

36. Sefiani, M., Le Caer, J. P., and Soyez, D. (1996) Characterization of hyperglycemic and molt-inhibiting activity from sinus glands of the penaeid shrimp Penaeus vannamei. Gen Comp Endocrinol 103, 41–53

37. Cooke, I. M., Sullivan,R.E. (1982) Hormones and neurosecretion. In: The Biology of Crustacea. H. Atwood and D. Sandeman, eds. Academic Press, New York 3, 206–290

38. Sommer, M. J., and Mantel, L. H. (1988) Effect of dopamine, cyclic AMP, and pericardial organs on sodium uptake and Na/K‐ ATPase activity in gills of the green crab Carcinus maenas (L). Journal of Experimental Zoology Part A: Ecological Genetics and Physiology 248, 272–277

39. Serrano, L., Blanvillain, G., Soyez, D., Charmantier, G., Grousset, E., Aujoulat, F., and Spanings-Pierrot, C. (2003) Putative involvement of crustacean hyperglycemic hormone isoforms in the neuroendocrine mediation of osmoregulation in the crayfish Astacus leptodactylus. J Exp Biol 206, 979–988

40. Charmantier-Daures, M., Charmantier, G., Janssen, K. P., Aiken, D. E., and van Herp, F. (1994) Involvement of eyestalk factors in the neuroendocrine control of osmoregulation in adult American lobster Homarus americanus. Gen Comp Endocrinol 94, 281–293

41. Spanings-Pierrot, C., Soyez, D., Van Herp, F., Gompel, M., Skaret, G., Grousset, E., and Charmantier, G. (2000) Involvement of crustacean hyperglycemic hormone in the control of gill ion transport in the crab Pachygrapsus marmoratus. Gen Comp Endocrinol 119, 340–350

42. Prymaczok, N. C., Pasqualino, V. M., Viau, V. E., Rodriguez, E. M., and Medesani, D. A. (2016) Involvement of the crustacean hyperglycemic hormone (CHH) in the physiological compensation of the freshwater crayfish Cherax quadricarinatus to low temperature and high salinity stress. Journal of comparative physiology. B, Biochemical, systemic, and environmental physiology 186, 181–191

43. Chung, J. S., Dircksen, H., and Webster, S. G. (1999) A remarkable, precisely timed release of hyperglycemic hormone from endocrine cells in the gut is associated with ecdysis in the crab Carcinus maenas. Proc Natl Acad Sci U S A 96, 13103–13107

44. Chung, J. S., and Webster, S. G. (2006) Binding sites of crustacean hyperglycemic hormone and its second messengers on gills and hindgut of the green shore crab, Carcinus maenas: a possible osmoregulatory role. Gen Comp Endocrinol 147, 206–213

45. Turner, L. M., Webster, S. G., and Morris, S. (2013) Roles of crustacean hyperglycaemic hormone in ionic and metabolic homeostasis in the Christmas Island blue crab, Discoplax celeste. J Exp Biol 216, 1191–1201

46. Sun, D., Lv, J., Gao, B., Liu, P., and Li, J. (2019) Crustacean hyperglycemic hormone of Portunus trituberculatus: evidence of alternative splicing and potential roles in osmoregulation. Cell Stress Chaperones 24, 517–525

47. Mantel, L. H., and Farmer, L. L. (1983) Osmotic and ionic regulation. Internal anatomy and physiological regulation 5, 53–161

48. Davenport, J., and Wong, T. (1987) Responses of adult mud crabs (Scylla serrata)(Forskal) to salinity and low oxygen tension. Comparative Biochemistry and Physiology Part A: Physiology 86, 43–47

49. Chen, J.-C., and Chia, P.-G. (1997) Osmotic and ionic concentrations of Scylla serrata (Forskål) subjected to different salinity levels. Comparative Biochemistry and Physiology Part A: Physiology 117, 239–244

50. Chou, P. Y. (1978) Prediction of the secondary structure of proteins from their amino acid sequence. Advances in enzymology and related areas of molecular biology 47, 45–148

51. Katayama, H., Ohira, T., Aida, K., and Nagasawa, H. (2002) Significance of a carboxyl-terminal amide moiety in the folding and biological activity of crustacean hyperglycemic hormone. Peptides 23, 1537–1546

52. Mosco, A., Edomi, P., Guarnaccia, C., Lorenzon, S., Pongor, S., Ferrero, E. A., and Giulianini, P. G. (2008) Functional aspects of cHH C-terminal amidation in crayfish species. Regul Pept 147, 88–95

53. Webster, S. (1993) High-affinity binding of putative moult-inhibiting hormone (MIH) and crustacean hyperglycaemic hormone (CHH) to membrane-bound receptors on the Y-organ of the shore crab Carcinus maenus. Proceedings of the Royal Society of London. Series B: Biological Sciences 251, 53–59

54. Katayama, H., and Chung, J. S. (2009) The specific binding sites of eyestalk- and pericardial organ-crustacean hyperglycaemic hormones (CHHs) in multiple tissues of the blue crab, Callinectes sapidus. J Exp Biol 212, 542–549

55. Chen, H.-Y., Toullec, J.-Y., and Lee, C.-Y. (2020) The crustacean hyperglycemic hormone superfamily: progress made in the past decade. Frontiers in Endocrinology 11, 578958.

56. Chung, K. F., and Lin, H. C. (2006) Osmoregulation and Na,K-ATPase expression in osmoregulatory organs of Scylla paramamosain. Comp Biochem Physiol A Mol Integr Physiol 144, 48–57

57. Chen, H. Y., Watson, R. D., Chen, J. C., Liu, H. F., and Lee, C. Y. (2007) Molecular characterization and gene expression pattern of two putative molt-inhibiting hormones from Litopenaeus vannamei. Gen Comp Endocrinol 151, 72–81

58. Schägger, H., and Von Jagow, G. (1987) Tricine-sodium dodecyl sulfate-polyacrylamide gel electrophoresis for the separation of proteins in the range from 1 to 100 kDa. Analytical biochemistry 166, 368–379

59. Zhou, H. X., Lyu, P., Wemmer, D. E., and Kallenbach, N. R. (1994) Alpha helix capping in synthetic model peptides by reciprocal side chain–main chain interactions: evidence for an N terminal “capping box”. Proteins: Structure, Function, and Bioinformatics 18, 1–7

60. Lobley, A., Whitmore, L., and Wallace, B. (2002) DICHROWEB: an interactive website for the analysis of protein secondary structure from circular dichroism spectra. Bioinformatics 18, 211–212

61. Peterson, G. L. (1978) A simplified method for analysis of inorganic phosphate in the presence of interfering substances. Analytical Biochemistry 84, 164–172

62. Holliday, C. W. (1985) Salinity‐ induced changes in gill Na, K‐ ATPase activity in the mud fiddler crab, Uca pugnax. Journal of Experimental Zoology Part A: Ecological Genetics and Physiology 233, 199–208

63. Tsai, J. R., and Lin, H. C. (2007) V-type H+-ATPase and Na+,K+-ATPase in the gills of 13 euryhaline crabs during salinity acclimation. J Exp Biol 210, 620–627

64. Golovanov, A. P., Hautbergue, G. M., Wilson, S. A., and Lian, L. Y. (2004) A simple method for improving protein solubility and long-term stability. J Am Chem Soc 126, 8933–8939

65. Goddard, T. D., and Kneller, D. G. (2008) SPARKY 3. University of California, San Francisco

66. Brunger, A. T. (2007) Version 1.2 of the Crystallography and NMR system. Nature protocols 2, 2728

67. Oostenbrink, C., Villa, A., Mark, A. E., and van Gunsteren, W. F. (2004) A biomolecular force field based on the free enthalpy of hydration and solvation: the GROMOS force-field parameter sets 53A5 and 53A6. Journal of computational chemistry 25, 1656–1676

68. Laskowski, R. A., MacArthur, M. W., Moss, D. S., and Thornton, J. M. (1993) PROCHECK: a program to check the stereochemical quality of protein structures. Journal of applied crystallography 26, 283–291

